# Analysis of variability of functionals of recombinant protein production trajectories based on limited data

**DOI:** 10.1101/2021.04.04.438346

**Authors:** Shuting Liao, Kantharakorn Macharoen, Karen A. McDonald, Somen Nandi, Debashis Paul

**Affiliations:** Graduate Group in Biostatistics, University of California, Davis, CA 95616, USA; Department of Chemical Engineering, University of California, Davis, CA 95616, USA; Global HealthShare, University of California, Davis, CA 95616, USA; Department of Statistics, University of California, Davis, CA 95616, USA

## Abstract

We propose a method for analyzing the variability in smooth, possibly nonlinear, functionals associated with a set of product production trajectories measured under different experimental conditions. The key challenge is to make meaningful inference on these parameters across different experimental conditions when only a limited number of measurements in time are collected for each treatment, and when there are only a few, or no, replicates available. For this purpose we adopt a modeling approach by representing the production trajectories in a B-spline basis, and develop a bootstrap-based inference procedure for the parameters of interest, also accounting for multiple comparisons. The methodology is applied to study two types of quantities of interest – “time to harvest” and “maximal productivity” in the context of an experiment on the production of certain recombinant proteins under laboratory conditions. We complement the findings by extensive numerical experiments that look into the effects of different types of bootstrap procedures and associated schemes for computing *p*-values for tests of hypotheses.

## Introduction

Many biological experiments involve measuring the production of certain biochemical compounds over a period of time under various experimental conditions. So, the data associated with such experiments are inherently longitudinal. One long-studied problem is to compare these production, (or growth) trajectories across different treatments, which is a core topic of longitudinal data analysis [1–4]. In most of these studies, the object of interest is typically the expected amount of the substance being measured, and one has multiple replicates to accommodate a comprehensive ANOVA (Analysis of Variance) approach to deal with the problem of ascribing treatment effects. Indeed, the traditional approach to such inferential questions has been through the application of repeated measures designs [5–7].

However, in many real-life laboratory experiments, one key constraint is the number of data points or replicates that can be obtained, due to the time and cost associated with completing the experiment. Furthermore, in many instances, as we discuss below, the key object of interest is not the level of the chemical substance itself, but some, possibly nonlinear, functionals of the production trajectory. For instance, this functional could be (a) the time it takes for the compound concentration to reach a prespecified value (to be referred to as time-to-harvest); (b) the maximum production level (= maximum of the production trajectory); (c) the maximum productivity, defined as the maximum of the amount divided by time over the duration of the experiment.

### Statistical challenge

One key requirement of the analysis of the variability of such functionals across different experimental conditions (treatments) is to ensure that the underlying production trajectories are monotonic, without which some of the quantities of interest are not properly defined. At the same time, due to the limited number of time points at which these production trajectories are typically measured, it is imperative to borrow information from the commonality of the structures in order to ensure that we have enough degrees of freedom left for comparing the functionals across the treatments. The other challenge is that, due to the limited number of data points, any inference procedure that directly relies on the asymptotic theory of the statistics involved, will have limited accuracy, or may be misleading. Moreover, since some of the parameters (functionals of the production trajectories), or process metrics, of interest are nonlinear, standard ANOVA framework that relies on linear models theory does not apply immediately. In this paper, we primarily focus on comparing the equality of the parameters across treatments, by means of simultaneous pairwise comparisons. So, the statistical challenge is to develop a methodology that ensures (a) monotonicity of the fitted trajectories; and (b) can handle simultaneous inference for arbitrary continuous functionals of the trajectories with a limited amount of data.

### Statistical methodology and key contributions

In order to address the challenges in terms of statistical inference described above, we adopt the following three-pronged approach. First, following the ideas in [8], we model the production trajectories by representing them in a B-splines basis with a fixed number of knots, and incorporate the monotonicity constraint by imposing linear inequality constraints on the B-spline coefficients. Our proposal furthermore allows for ascribing the variability in the shape of the production trajectories to the treatment and temporal effects and their potential interactions, though we do not pursue this direction much further since comparing the entire trajectories across treatments is not the main focus of this work. We fit the trajectories by using a least squares regression procedure that is implemented through a quadratic programming approach.

Next, for statistical inference, we use bootstrap, or resampling procedure, as our main vehicle. We compare two different versions of bootstrap, namely, the residual bootstrap and parametric bootstrap. Both these methods are well-adapted to deal with the problem of comparing the values of parameters that are arbitrary functionals of the production trajectories corresponding to different treatments.

Finally, since we are pursuing simultaneous inference involving many pairwise comparisons, we adopt a method for imparting control on the false discovery rate while constructing the simultaneous confidence intervals for the pairwise differences, by pursuing a technique presented in [9]. In summary, we provide a comprehensive framework for simultaneous statistical inference on several process metrics that are functionals of biochemical production (or growth) trajectories, based on fairly limited amount of data, with empirical validity.

## Scientific context

Butyrylcholinesterase (BChE) circulating in human blood plasma is a tetrameric hydrolase enzyme that can be potentially used as a prophylactic and/or therapeutic treatment against organophosphorus nerve agents [10]. However, the use of purified BChE from human blood plasma in clinical stage is limited by its cost that was estimated to be $20,000 per 400-mg dose [11]. Thus, recombinant human BChE (rBChE) has been developed in several host expression systems, including transgenic rice cell suspension cultures, to be used as an alternative source of BChE. Our lab has developed metabolically regulated transgenic rice cell suspensions under the RAmy3D promoter to produce rice-made recombinant human BChE (rrBChE) [12, 13]. In nature, the RAmy3D promoter in rice cells derived from rice seed, is suppressed in sugar-rich environment but activated in sugar-starved environment [14–16]. In other words, the RAmy3D promoter-based transgenic rice cell suspensions are grown is sugar-rich medium for production and transferred into sugar-free medium for rrBChE production.

A major challenge of growing plant cell suspension cultures is the slow growth rate of plant cells compared to microbial and mammalian cells. For example, it takes 6–7 days for transgenic rice cells to reach mid-to-late exponential growth phase, following by the medium exchange to replace spent growth medium with sugar-free medium, and another 4–5 days post induction for rrBChE expression [12, 13]. In other words, the cultivation time of transgenic rice cell suspensions in a batch culture is 10–12 days. When it comes to an experiment with several treatments within time and equipment constraints, the number of bioreactor replicates is likely to be restricted due to time of cultivation and limited equipment. Therefore, in this study, we employ novel statistical approaches to tackle limited data to build trajectory models using previously reported bioreactor data [13] to predict interested outcomes, such as time-to-harvest, maximum rrBChE production level, maximum productivity, and early stopping time. In addition, estimating the trajectory of the production level is a part of Quality by Design (QbD) [17] that is essential in biomanufacturing where a computationally feasible statistical method is involved in modeling based on available data.

## Data collection method

A 5L bioreactor (BioFlo 3000, formerly New Brunswick Scientific, Eppendorf Inc., Hauppauge, NY) was used to study the production of rrBChE under eight different conditions as previously described [13], and summarized in Table 1. In brief, the effects of dissolved oxygen (DO) were conducted in treatments (runs) A–E using a two-stage batch culture (the medium exchange was performed to replace spent sugar-rich medium with sugar-free medium to induce the promoter).

**Table 1.** Conditions used in bioreactor runs A–H when agitation rate and temperature were maintained at 75 rpm and 27^*o*^ C, respectively, in all runs. The aeration rate was maintained at 0.2 vvm (volume of sparged gas per working volume per minute) in runs A–F but 0.2–0.4 vvm in runs G and H (reproduced from [13])

| Experiment # | %DO during growth phase | %DO during induction phase | Media exchange |
| --- | --- | --- | --- |
| A | 40 | 10 | Yes |
| B | 40 | 20 | Yes |
| C | 40 | 30 | Yes |
| D | 40 | 40 | Yes |
| E | 40 | Uncontrolled | Yes |
| F | 40 | 40 | No |
| G | Uncontrolled | Uncontrolled | No |
| H<sup>1</sup> | Uncontrolled | Uncontrolled | No |

<sup>1</sup> Initial sucrose concentration in the medium in run H was reduced to 15 g/L instead of 30 g/L used in other runs. DO, dissolved oxygen.

Treatments F–H were operated in single-stage batch culture (no medium exchange; production was simply induced by sugar depletion from cellular uptake) with or without controlling DO and using 50% of the usual initial sucrose concentration during the growth phase. For each treatment, samples were taken every day during days 0–5 post induction (dpi) to quantify rrBChE activity in cell extract and culture medium using the modified Ellman assay [18], and assuming a specific activity of 260 U/mg crude rrBChE to convert activity to rrBChE production level (ug/g fresh weight of rice cells) [13].

## Scientific goals of the study

One of the goals of this study is to analyze variability of production trajectories of limited data set of the recombinant protein production by using rrBChE as a model study. Another goal is to use statistical approaches to determine an optimal time to harvest a recombinant protein during a protein production process.

## Statistical methodology

### Modeling production trajectories

We suppose there are *I* ≥ 2 treatments and each treatment is applied to several independently chosen experimental units (bioreactors). Further, the response (e.g. rrBChE concentration in the bioreactor) is measured at observation times 0 < *t*_*i*1_ < … < *t*_*iJ*_ = *T*, say, for *J* ≥ 2 (this allows the observation times to be different for different treatments). Let us denote the mean response curve (at time *t* ≥ 0) corresponding to the *i*-th treatment as *µ*_*i*_(·). It is assumed that *µ*_*i*_(·) is a monotonically increasing function of time, over the observation time window [0, *T*].

For simplicity as well as statistical efficiency, we assume a balanced design, that is, the sample size at each observational time is the same for every treatment. The measurement process is destructive, so that for any particular experimental unit, we only have one measurement, at the time of sampling the bioreactor. Therefore, to obtain reasonably accurate measurement for the whole trajectory, we need replicates, for each time *t*_*ij*_ and each treatment *i*. Let *n* denote the number of replicates assigned to each combination (*i, j*), which corresponds to a balanced design. Note that we allow *n* = 1 since in practice only limited data is available especially in some biological experiments. Denote the response from the *k*-th experimental unit, in the *i*-th treatment group, measured at time *t*_*ij*_, to be *Y*_*ijk*_.

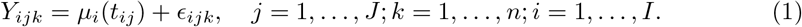

where *ϵ*_*ijk*_ are independent random variables with mean 0 and unknown, common variance *σ*^2^ *>* 0.

In practice, we may allow the number of time points *J* to depend on index *i* as well. We use a basis representation approach for modeling the mean trajectories *µ*_*i*_’s. In particular, we adopt cubic B-spline basis functions [19] for representing the functions. For each *i*, cubic B-spline basis functions are used (assuming *L* cubic B-spline basis) to model *µ*_*i*_(*t*):

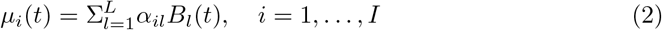

One advantage of spline representation is that, since the functions *B*_*l*_(·)’s are non-negative, the curves *µ*_*i*_’s are nonnegative provided the coefficients *α*_*il*_’s are so. A more significant advantage, from the point of view of modeling “production curves” of the type considered here, is that the condition *µ*_*i*_(*t*) is non-decreasing in *t* can be imposed by simply requiring that

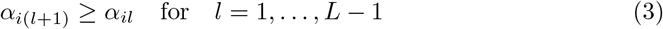

for all *i* [8, 20]. This is a reasonable assumption for batch production of a recombinant protein if the protein is stable in the culture medium (e.g. there is no consumption and/or degradation of the product, simply accumulation due to production). In addition, we incorporate the treatment effects and treatment-time interaction by modeling *α*_*il*_’s as follows:

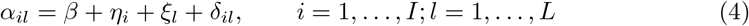

For identifiability purposes, we impose the following restrictions:

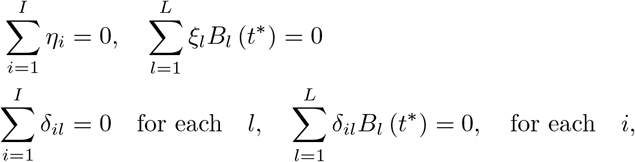

where *t*^*\**^ is an arbitrary but appropriately chosen point in [0, *T*]. And the constraints Eq (3) are equivalent to

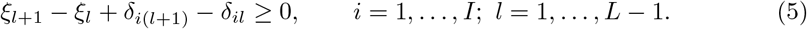

The model given by Eq (1), Eq (2) and Eq (4) is a modified form of two-factor ANOVA and can be solved by least square procedures under identifiability conditions. Assuming that the model specification is correct, the resulting estimate of *µ*_*i*_(*t*) for any *t* will be unbiased and will have an approximate normal distribution for reasonably large values of *n*. In this case, we can rely on large sample theory for statistical inference on the parameters of interest.

However, in the applications considered here, we need to take into account the monotonicity constraints Eq (3) (or Eq (5)) for modeling the production/growth trajectories. The least square procedures for fitting the model given by Eq (1), Eq (2), Eq (4) and Eq (5) results in a **quadratic programming problem**. Though such estimates guarantee the monotonicity of the mean response curve, the estimates of *µ*_*i*_(*t*)’s are not unbiased. The biases are small but non-negligible. The number *L* of basis functions used to model the growth/production trajectories is a user defined quantity that controls the degree of complexity of the trajectories, with larger values allowing for more complex shapes. Larger *L* also reduces the degree of bias, at the cost of increasing the variance of the fits. In practice *L* may be determined either in an ad-hoc manner, or by utilizing data from pilot studies, through a cross-validated curve-fitting procedure. Usage of monotonicity constrained estimators and limited number of replicates both limit the application of classical large sample theory in dealing with the inference problem. Instead, we focus attention on resampling-based approached for solving the problem of statistical inference.

## Inferential questions

We present the mathematical formulation of the inferential questions associated with the parameters of interest mentioned earlier.

### Time-to-harvest

*θ*_*i*_ = min {*t* : *µ*_*i*_(*t*) = *c*} for *i* = 1, …, *I*, where *c* is the prespecified cut-off level. The corresponding null hypothesis representing no treatment effect on the time-to-harvest is:

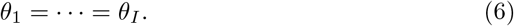

With 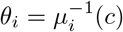 for some given cut-off level *c*, we are interested in testing the one-sided null hypotheses of the form *θ*_1_ ≥ *s*_1_, …, *θ*_*I*_ ≥ *s*_*I*_ (here the times *s*_1_, …, *s*_*I*_ need not be equal). These hypotheses translate to the linear inequality constraints:

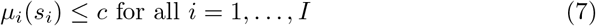

Notice that, *θ*_1_ = · · · = *θ*_*I*_ is not a linear constraint.

- However, the null hypothesis *θ*_1_ =· · · = *θ*_*I*_ = *s*_0_ can be translated to the equality constraints:

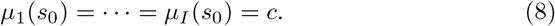

- A composite null hypothesis of the form *s*_*L*_ ≤ *θ*_*i*_ ≤ *s*_*U*_ for all *i* can also be translated into linear inequality constraints

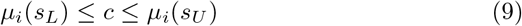

for all *i*.

### Maximum production

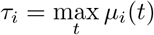 for *i* = 1, …, *I*. The regarding null hypotheses are

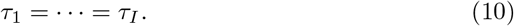

or equivalently

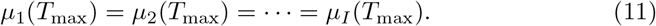

where *T*_max_ is the largest time point during the experiment.

### Maximum “unweighted” productivity

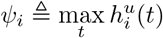, for *i* = 1, …, *I*, where 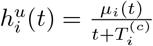 represents “unweighted” productivity of the *i*-th treatment and 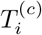 is the number of days of cultivation before the induction. The corresponding null hypotheses are

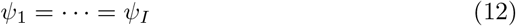

or equivalently

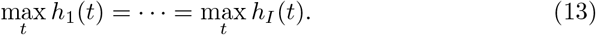

### Early stopping time

Suppose the decision to harvest is taken based on the relative gradient of *µ*_*i*_(*t*) (or, gradient of log *µ*_*i*_(*t*)). Let 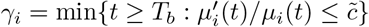 where *T*_*b*_ is some constant baseline time and 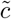 is a gradient threshold. We may be interested in testing hypotheses of the form *γ*_*i*_ *> s*_0_ for *i* = 1, …, *I* where time *s*_0_ is treated as the same for all i for simplicity.

Since 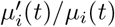 is not necessarily monotone, this cannot be easily reduced to a simple set of inequality constraints. However, we may discretize time to a grid of the form *T*_*b*_ = *T*_1_ <· · · < *T*_*m*_ = *s*_0_ and then consider the slightly relaxed form of the hypothesis

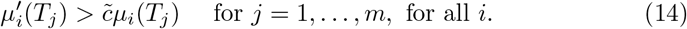

In practice, *m* needs to be pretty small for the feasibility of the optimization problem.

In a more general sense, if our interest is in testing for equality of Θ_*i*_’s where Θ_*i*_ is a linear functional of *µ*_*i*_(*t*), then we can design a procedure that imposes a less restrictive null hypothesis. This applies, for example, to the case when the maximum production level is the quantity of interest, since under monotonicity of *µ*_*i*_, this is simply the value at the maximum time point.

Specifically, suppose Θ_*i*_ = *A*(*µ*_*i*_) where *A* is a linear operator taking a scalar value. Then Θ_*i*_ can be expressed as Σ _*l*_ *r*_*l*_*α*_*il*_ where *r*_*l*_ are some known coefficients and *α*_*il*_ represents the *l*-th spline coefficient of *µ*_*i*_(*t*) =Σ_*l*_ *α*_*il*_*B*_*l*_(*t*).

- Example 1: if Θ_*i*_ = *µ*_*i*_(*T*), then *r*_*l*_ = *B*_*l*_(*T*).
- Example 2: if Θ_*i*_ = *µ*_*t*_(*t*)*dt*, then *r*_*l*_ = *B*_*l*_(*t*)*dt*.

## Inference using bootstrap

As we mentioned before, 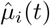 estimated by imposing constraints Eq (3) or Eq (5) is biased. What matters more is that we have such limited number of replicates that we cannot trust Large Sample Theory. In view of this, it is imperative to adopt an inferential framework that does not depend too heavily either on the model assumptions, or the methodology, or indeed the sample sizes. Therefore, as a possible alternative, we propose to infer about Θ_*i*_’s which represent any parameters of interest we mentioned above, by making use of a non-parametric (or parametric) bootstrap procedure.

### Confidence interval for one parameter

When the number of replicates *n* is relatively but not extremely small, we construct the confidence interval for one parameter (e.g., the difference between a fixed pair of treatments) as follows, using a nonparametric bootstrap procedure. We re-sample with replacement randomly, samples of size *n*, from the collection of observations 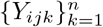 for each treatment-time pair (*i, j*). We repeat this process B times and denote 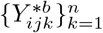 as the *b*-th sample corresponding to pair (*i, j*). Then we fit the model given by Eq (1), Eq (2), Eq (4) and Eq (5) and obtain a set of estimated parameters we are interested, denoted by 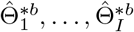.

As an alternative, one can use parametric bootstrap method to obtain bootstrap estimates, if there is a high degree of confidence in the model assumption or the number of replicates is very small. The procedures for parametric bootstrap method are as follows:

- Calculate the sample variance for the *i*-th treatment at time *t*_*j*_ based on *n*_*i*_ replicates (for balanced data: *n*_*i*_ = *n*): 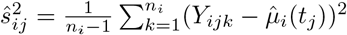
- Compute the pooled variance 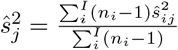
- Simulate parametric bootstrap samples 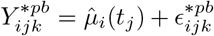 where 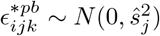.

We propose two methods to construct the bootstrap confidence intervals:

1. **Percentile bootstrap confidence intervals**: We obtain percentile bootstrap confidence intervals for Θ_*i*_ and Θ_*i*_ − Θ_*j*_ by *B* estimates of these parameters. The intervals will depend on appropriate quantiles of the bootstrap estimates 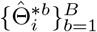 and 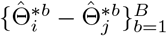, respectively.
2. **Bias-corrected and accelerated bootstrap interval** *BC*_*a*_: Percentile bootstrap confidence interval is only first-order accurate. In addition, it can be inaccurate for skewed distribution, that is, it does not correct for skewness. When the bootstrap distributions for some parameters are skewed, we propose to use bias-corrected and accelerated bootstrap intervals efron1987, denoted by *BC*_*a*_, which is not only second-order accurate but also corrects for the skewness in the sampling distribution.

## *p*-value associated with tests of hypothesis

Computation of the *p*-values is an essential part of the inference procedure, especially so when we perform simultaneous hypotheses testing. We propose two ways of obtaining approximate *p*-values using bootstrap. One is the percentile(-*t*) bootstrap, a general procedure, while the other one is a special case where the sample distribution of the test statistics under the null hypothesis can be calculated. For a general description different types of bootstrap procedures and their theoretical validity, one may refer to [22].

## *p*-value computation by percentile-*t* bootstrap

The idea is to use the duality between hypothesis testing and confidence intervals. For example, suppose we are performing pairwise tests for equality of the Θ_*i*_’s, where Θ_*i*_ is some functional of *µ*_*i*_, by doing percentile-*t* bootstrap. The 100(1 − *α*)% bootstrap confidence intervals for *δ*_*ij*_ = Θ_*i*_ − Θ_*j*_ are of the form

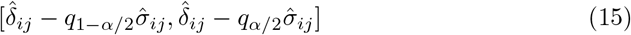

where 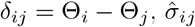 is the estimated standard error of 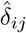 and *q*_*p*_ is the *p*-th quantile of the bootstrap distribution of the *t*-statistics

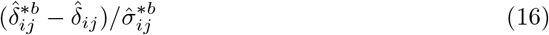

where 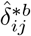 is the bootstrap estimate of *δ*_*ij*_ and 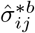 is the bootstrap estimate of its standard error. Then, we may define a bootstrap *p*-value for testing 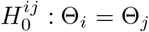 to be the largest value of *α* such that the aforementioned 100(1 −*α*)% bootstrap confidence interval contains 0. Note that 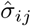 and 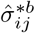 can be estimated by using “double bootstrap” procedure described in the S1 Appendix.

## *p-*value computation by percentile bootstrap

Alternatively, we may use the percentile bootstrap (arguably less accurate) instead of percentile-*t* bootstrap, to compute the *p*-values. The percentile bootstrap is less computationally intensive since it does not require computation of 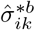 for each bootstrap sample, and so there is no need for the second layer of bootstrap. We only need to find the bootstrap *p*-value to be the largest value of *α* such that the 100(1 − *α*)% bootstrap confidence interval contains 0.

## Special case – null distribution through bootstrap

In some instances, the sampling distribution of the test statistics under the null hypothesis can be calculated through a bootstrap procedure. Then there is a different way of obtaining approximate *p*-values using bootstrap.

In each of the aforementioned hypothesis testing problems, the estimation problem under the null hypothesis remains a quadratic programming problem, with linear equality or inequality constraints on the parameters. In each case, the key parameters involved can be expressed in the form

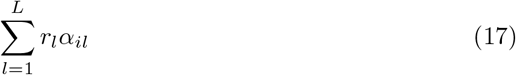

where *r*_*l*_’s are some constants and *α*_*il*_ is the coefficient of *µ*_*i*_(*t*) associated with the *l*-th B-spline *B*_*l*_(*t*). Since the estimation problem is a quadratic programming problem, we can use the fitted trajectories to construct surrogate data and then make use of this to compute the null distribution of the test statistics, and thereby obtain the *p*-value of the test. The surrogate data are computed as

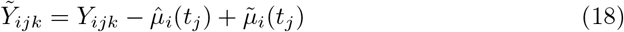

where 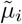 is the estimate of *µ*_*i*_ under the constraints imposed by the null hypothesis.

Here are the steps of the bootstrap procedure for approximating the sampling distribution under the null hypothesis of no difference across the different treatments:

1. Use the current method to compute 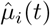 for the different treatments.
2. Obtain the fitted curves 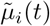 for the different treatments from the original data by imposing the additional constraint on the coefficients (*α*_*il*_) that

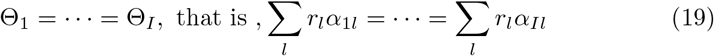
3. Create surrogate data 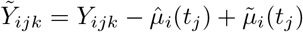, *i* = 1, …, *I, j* = 1, …, *J*. 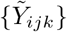 are under the null hypothesis of no treatment effect.
4. Apply the current procedure for generating the parametric or non-parametric bootstrap sampling distribution to the surrogate data 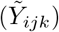 to generate bootstrap estimates 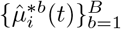. This leads us to a set of bootstrap estimates of Θ_*i*_,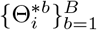
5. Use the bootstrap sampling distribution computed in Step 4 to obtain the *p*-values for testing Θ_*i*_ = Θ_*i′*_ for all pairs (*i, i*^*′*^).

Notice that Step 2 of the above procedure imposes linear equality constraints on the parameters (*α*_*il*_), along with the original monotonicity and non-negativity constraints, and therefore the resulting least squares problem can still be solved by quadratic programming. Thus, Step 2 obtains least squares estimates of *µ*_1_,· · ·, *µ*_*I*_, under the monotonicity constraint and the additional requirement (null hypothesis) that Θ_1_ = · · · = Θ_*I*_. This is clearly less stringent than requiring that *µ*_*i*_’s are all equal. In step 5, for example, if 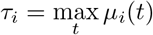, then the *p*-value for the test will be computed as the fraction of times 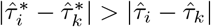. Here, 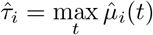 (with 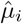 computed in Step 1) and 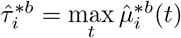 (with 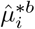 computed in Step 4).

In spite of the simplicity of this method, if the null hypotheses is not be in a form that enables generation of surrogate data from the null distribution, we can not use the method described under the “Special case”. Instead, in such cases we make use of percentile-*t* bootstrap or percentile bootstrap. It is interesting that in the cases where we have alternate ways of computing the bootstrap *p*-values, the two versions of *p*-values are similar in the particular real data analysis that we performed.

## Adjusted confidence intervals and *p*-values for simultaneous inference

If we are to perform tests for multiple hypotheses of the kind *H*_0_ : Θ_*i*_ = Θ_*i′*_ vs. *H*_1_ : Θ_*i*_ = Θ_*i*_ for several pairs of treatments 1 ≤ *i* < *i*^*′*^ ≤ *I*, then we can perform simultaneous hypotheses tests (equivalently, obtain simultaneous confidence intervals CI) by doing appropriate correction to the significance levels of each pairwise test to meet a prespecified level of familywise significance. This can be achieved by making use of the Bonferroni procedure or a False Discovery Rate (FDR) control procedure [23], where *p*-values need to be involved and calculated.

Once we obtain the bootstrap *p*-values (either by general procedure or the special case), the Benjamini-Hochberg (BH) procedure can be used for FDR correction to determine the significance of the different pairwise tests for a given level of significance. After that, we can adjust the confidence levels of the confidence intervals (FCR-Adjusted BH-Selected CIs) for the parameters accordingly [9]:

1. Sort the *p*-values used for testing the m hypotheses regarding the parameters, *p*_(1)_ ≤ · · · ≤ *p*_(*M*)_, where *M* is the number of tests.
2. Calculate 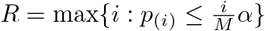 where *α* is the significance level.
3. Select *R* parameters for which 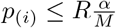, corresponding to the rejected hypothesis.
4. Construct a 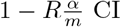 CI for each parameter selected.

## Two varieties of bootstrap

When the number of replicates is relatively small (e.g., *n* = 1), the bootstrap method is more reliable than the ones based on asymptotic theory. We primarily focus on two versions. One is the residual (nonparametric) bootstrap and the other is the parametric bootstrap.

### Residual bootstrap

The general route of generating residual bootstrap samples is as follows:

1. Suppose {*Y*_*ijk*_} as the raw data and obtain the 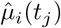 ;
2. Obtain 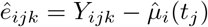, which is biased due to the framework;
3. For each *j*, obtain 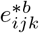 resampled from {*ê*_*ijk*_ − *ē*_.*j*._} for *i* = 1, …, *I* and *k* = 1, …, *n* where 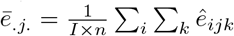. Note that {*ê*_*ijk*_ − *ē*_.*j*._} will sum to 0 for each *j*.
4. Generate residual bootstrap samples: 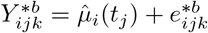.

One thing to note is that residual bootstrap is not effective especially when the number of treatment is small (say if *I* = 3, then the residual bootstrap method is a bad idea since resampling space is limited). This can be a practical challenge that one needs to make a choice of bootstrap procedures.

### Parametric bootstrap

One can also make use of parametric bootstrap as an alternative to generate bootstrap estimates when there are limited number of replicates and limited number of treatments. We come up with parametric bootstrap method by using the standard deviations 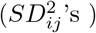. Note that here we make use of pooled variance 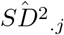 for all treatments at time *j* based on 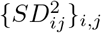. That is, 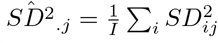. We use two different strategies to generate bootstrap samples. One is using normal distribution while the other one is *t*-distribution, which allows for more extreme values.

- **Gaussian noise:** At each time *b* = 1, …, *B*, we generate samples by 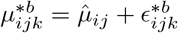, where 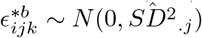. We use 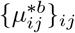 for model fitting, leading to 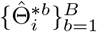.
- *t***-distributed noise:** Instead of assuming noise approximately follows normal distribution, we use *t*-distribution to allow more extreme values. We obtain the scale parameter by 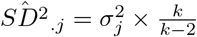 where *k* is the degree of freedom of *t*-distribution; 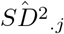 is the measurement error (pooled variance). Then we generate noise which follows *σ*_*j*_*t*_*k*_. Here we assume *k* = 3, which means the variance is finite.

In many instances, Gaussian assumption of noise could be too strong since we may only see values within 2 standard deviations. However, we should expect more extreme values in practice. Therefore, for the application to the rrBChE data, we use scaled *t*-distribution with relatively low degrees of freedom, which allows for extreme values. Using *t*-distributed noise indicates greater variability (Fig 1) and more extreme values of the bootstrap fitted curves, which reflects the variability in the real data more effectively. By comparing the confidence intervals, we can see the results in this real data application are similar regardless of the type of noise, affirming a degree of robustness of the method.

**Fig 1.**
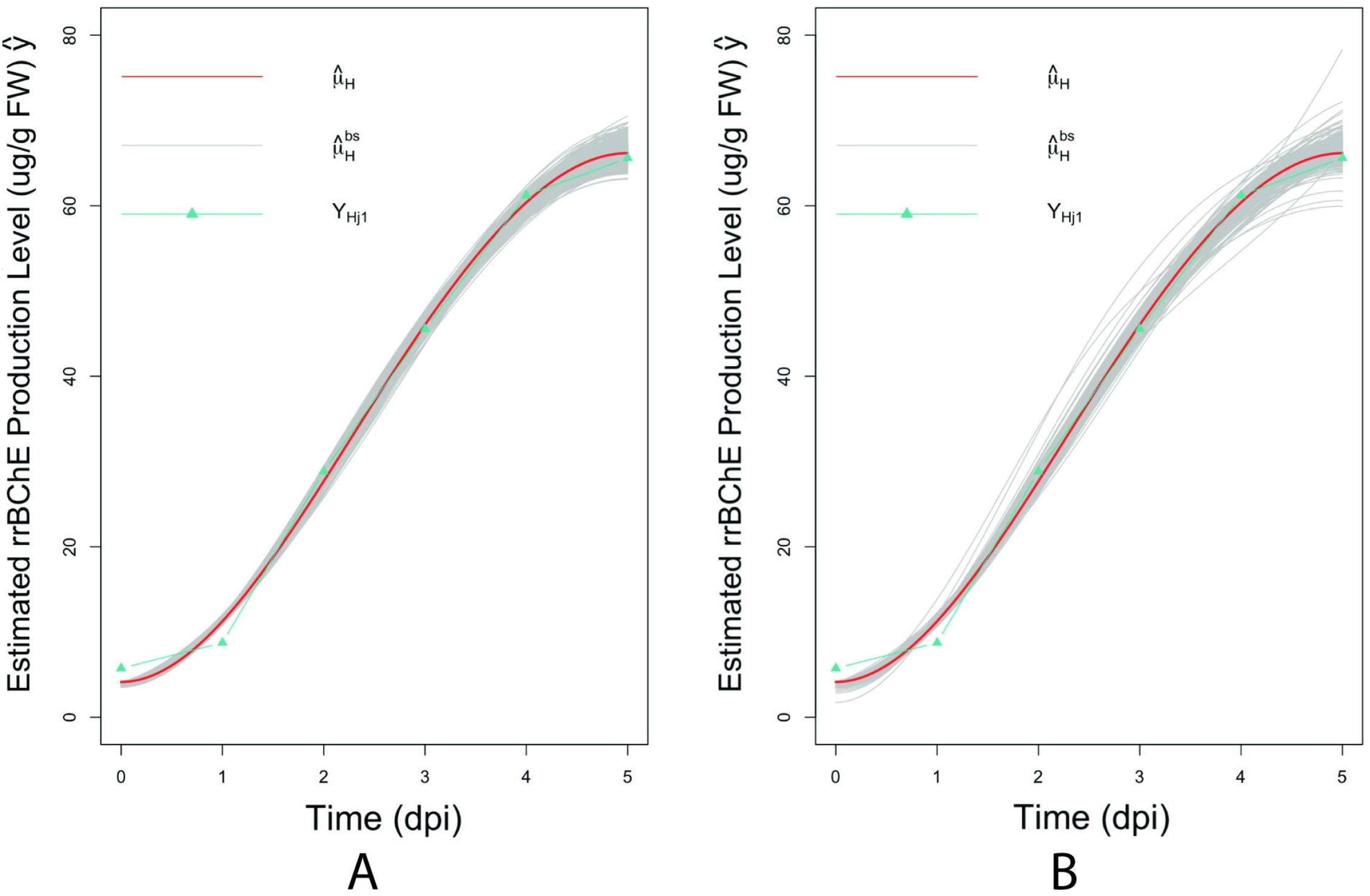
Observations, fitted curves and 500 bootstrap fitted curves for treatment H. A: assuming normal noise; B: assuming *t*-distributed noise.

## Results

### Simulation study

In this subsection, we present a simulation study illustrating the effectiveness of the proposed bootstrap-based inference procedures. This numerical simulation also allows us to make a comparison among the different bootstrap procedures.

#### Settings

We assume the number of treatments *I* = 3 and the time interval is (0, 9] with number of time points *J* = 9; All treatments share the same time points {1, 2, 3, 4, 5, 6, 7, 8, 9}. Assume there are *L* = 5 basis functions for cubic B-splines with equally spaced knots. For each time point, we have *n* = 5 replicates. Suppose *Y*_*ijk*_ = *µ*_*i*_(*t*_*ij*_) + *ϵ*_*ijk*_, where the noise level is 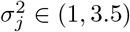. Our main interest is time-to-harvest *θ*_*i*_’s.

The simulated data and estimation by the standard least square procedure (without constraints) and quadratic programming framework (with constraints) are shown in Fig 2. Though the estimations using two methods are very similar in our case, there is difference. The estimated curve by least square procedure for Treatment 3 shows decreasing pattern by the end of time, while the one with constraints remains non-decreasing (see Fig A1 in S1 Appendix). Since we are interested in estimating the growth curve and ‘time-to-harvest’, how we fit the data really matters. In addition, using the standard least square procedure may result in oscillations in estimation. Suppose the time-to-harvest parameter *θ*_*i*_’s are of interest with a pre-specified level *c* = 10.3 (Fig 3) and we use the residual (nonparametric) bootstrap method for inference. We compute both percentile bootstrap confidence intervals and bias-corrected and accelerated bootstrap interval (*BC*_*a*_) since the bootstrap sampling distributions involving *θ*_2_ are skewed (Table 2 and Table 3). Both types of intervals suggest that *θ*_1_, *θ*_2_ are significantly different from *θ*_3_ respectively, though we can see *BC*_*a*_’s are slightly better than general ones in terms of the length of intervals.

**Table 2.** CI from Non-parametric Bootstrap for *c* = 10.3; (*θ*_1_, *θ*_3_) and (*θ*_2_, *θ*_3_) are significant pairs.

|  | CI lower | CI upper | CI length | true | mean | sd |
| --- | --- | --- | --- | --- | --- | --- |
| $\hat{\theta}_1^{bs}$ | 5.820 | 7.027 | 1.207 | 6.378 | 6.320 | 0.327 |
| $\hat{\theta}_2^{bs}$ | 5.892 | 9.000 | 3.108 | 7.387 | 7.739 | 0.995 |
| $\hat{\theta}_3^{bs}$ | 4.730 | 5.162 | 0.432 | 4.910 | 4.917 | 0.113 |
| $\hat{\theta}_1^{bs} - \hat{\theta}_2^{bs}$ | -3.063 | 0.568 | 3.631 | -1.009 | -1.418 | 1.046 |
| $\hat{\theta}_2^{bs} - \hat{\theta}_3^{bs}$ | 0.937 | 4.198 | 3.261 | 2.477 | 2.822 | 1.002 |
| $\hat{\theta}_3^{bs} - \hat{\theta}_1^{bs}$ | -2.171 | -0.820 | 1.351 | -1.468 | -1.403 | 0.350 |

**Table 3.** Bias-corrected and accelerated bootstrap interval (*BC*_*a*_) for *c* = 10.3; (*θ*_1_, *θ*_3_) and (*θ*_2_, *θ*_3_) are significant pairs.

|  | CI lower | CI upper | CI length |
| --- | --- | --- | --- |
| $\hat{\theta}_1^{bs}$ | 5.640 | 6.568 | 0.928 |
| $\hat{\theta}_2^{bs}$ | 6.036 | 9.000 | 2.964 |
| $\hat{\theta}_3^{bs}$ | 4.695 | 5.126 | 0.431 |
| $\hat{\theta}_1^{bs} - \hat{\theta}_2^{bs}$ | -3.189 | 0.108 | 3.297 |
| $\hat{\theta}_2^{bs} - \hat{\theta}_3^{bs}$ | 1.117 | 4.252 | 3.135 |
| $\hat{\theta}_3^{bs} - \hat{\theta}_1^{bs}$ | 1.766 | -0.525 | 1.241 |

**Fig 2.**
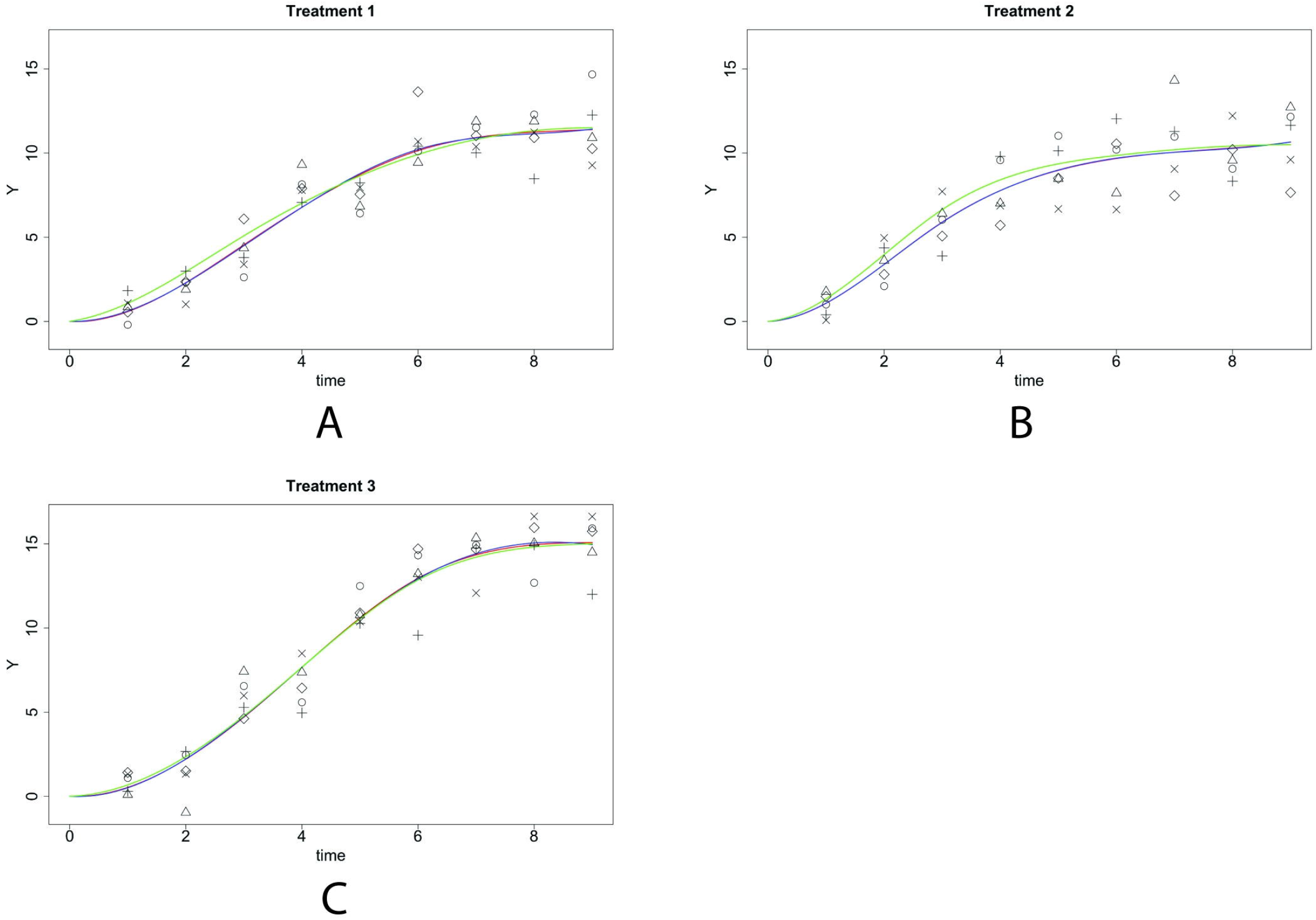
**Simulated data and estimation by the standard least square procedure and quadratic programming framework for different treatments; true data are denoted by different types of points; true curve *µ*(*t*) is marked in green; fitted curves with linear constraints are in red while the fitted curves by standard least square procedure are in blue.** A: Treatment 1; B: Treatment 2; C: Treatment 3.

**Fig 3.**
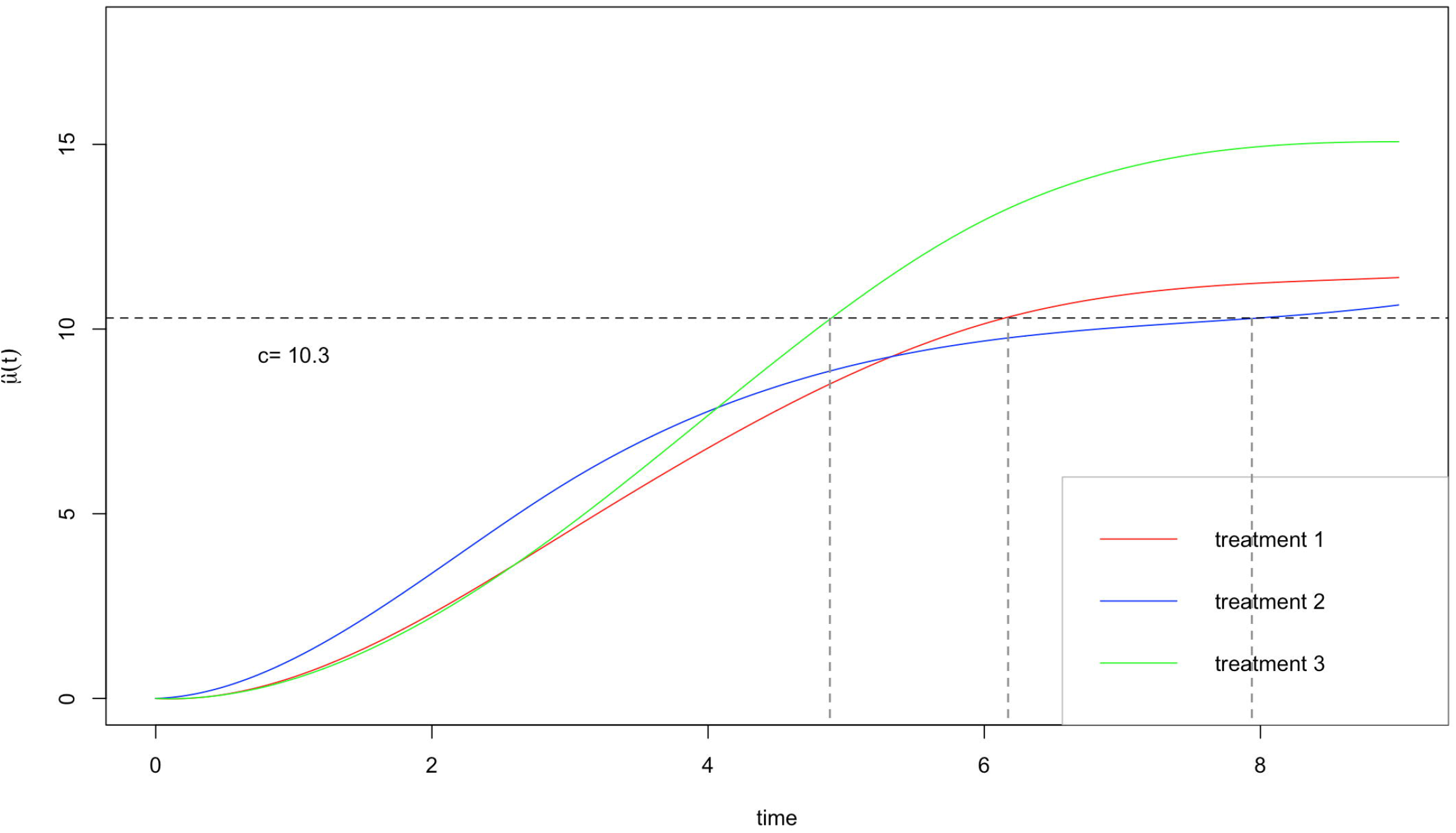
Fitted curves estimated with constraints; 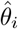 ‘s given by a specified level (*c* = 10.3) are indicated by vertical lines.

### Analysis of rrBChE data

For the rrBChE data, we have *I* = 8 treatments labeled A to H. We primarily focus on the protein production level (ug rrBChE/g FW rice cells) after sugar induction. We have *J* = 6 time points. Though one potential issue is that we only have one replicate (*n* = 1) at each time for each treatment, our framework is able to handle this and make appropriate inference. The observed data and estimation are shown in Fig 4.

**Fig 4.**
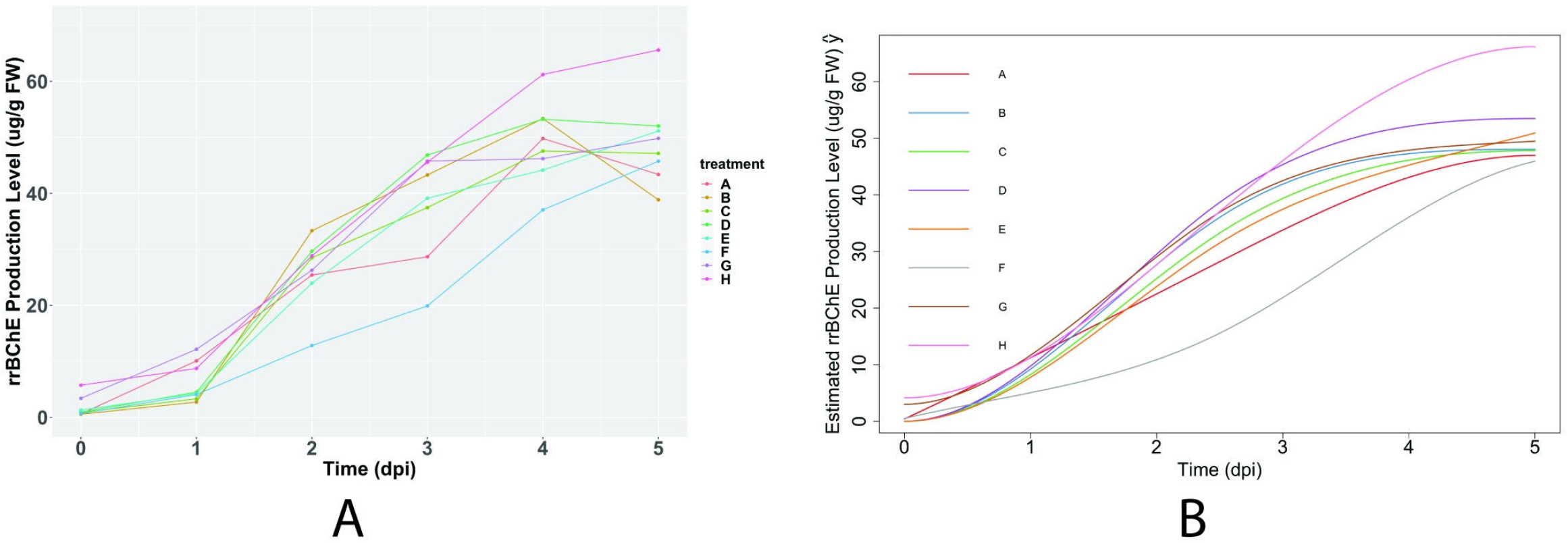
rrBChE data and the estimation. A: observed data; B: Fitted curves with monotonicity constraints.

#### Time-to-harvest *θ*_*i*_ and early stopping time *γ*_*i*_

With the pre-specified level *c* = 40 (ug/g FW), we obtain the estimates 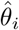’s and make inference by using the parametric bootstrap method. As for the early stopping time *γ*_*i*_, we set the level *c* = 0.1 and do the same things. We compute the corresponding confidence intervals by using normal noise and *t*-distributed noise separately. It turns out that the results are not sensitive to the type of noise (normal or *t*-distributed) we are using. The confidence intervals of *θ*_*i*_ and *γ*_*i*_ are shown in Fig 5. More details about the related confidence intervals can be found in Table A4 and Table A5 in S1 Appendix.

**Fig 5.**
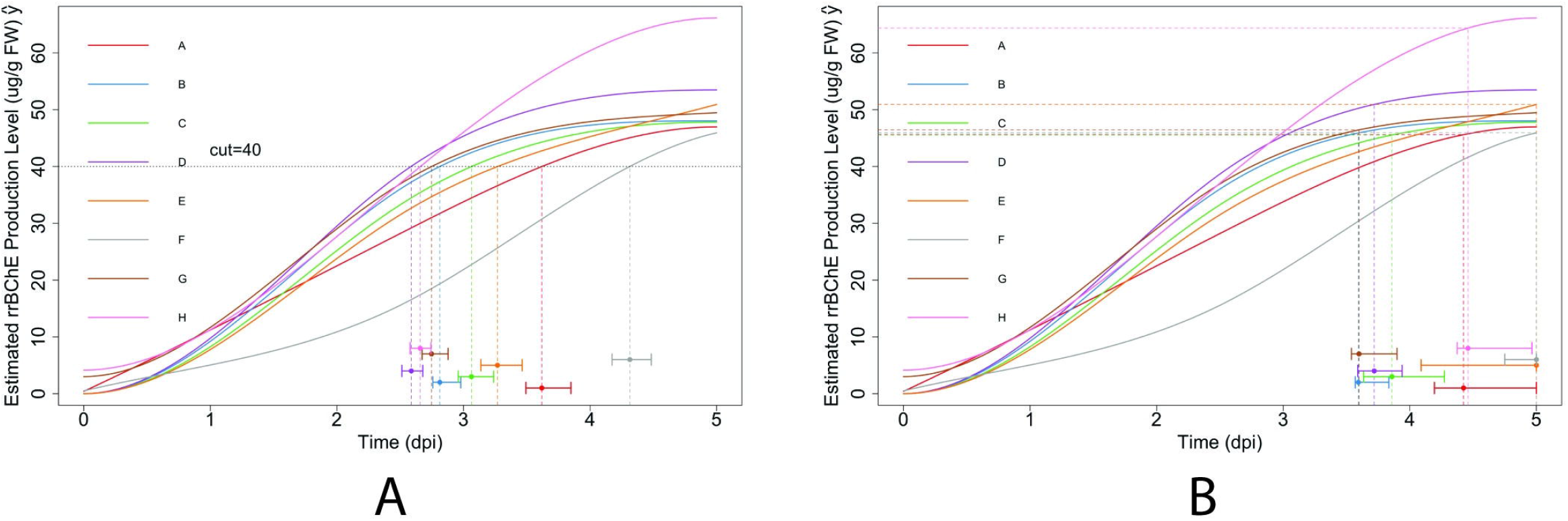
Estimated curves, 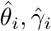 and their CIs, respectively. A: Fitted curves with specified level (cut-off *c* = 40); bootstrap confidence intervals of *θ*_*i*_’s are also shown (using *t*-distributed noise); B: Fitted curves with specified level 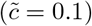 ; bootstrap confidence intervals of *γ*_*i*_’s are also shown (using *t*-distributed noise).

#### Simultaneous inference – maximum production level *τ*_*i*_

The hypotheses are:

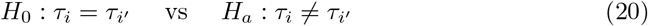

where 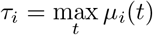 for *i* = 1, …, *I*. We use the Benjamini-Hochberg procedure for FDR control. Since the null hypothesis enables computation of estimates the quadratic programming framework, we use both nonparametric (residual) bootstrap and parametric bootstrap (using *t*-distributed noise) to compute two versions of *p*-values. It turns out that the rankings of *p*-values by the two different variants of bootstrap procedures are highly positively correlated, which means our method is robust and not sensitive to the way we compute *p*-values. All indicated significant pairs by residual bootstrap and percentile bootstrap CI method are shown in Table 4 and Table 5. The results related to using the null bootstrap distribution to compute *p*-values are shown in Table A1 and Table A2 in S1 Appendix. It is interesting to see that two versions of *p*-values indicate similar results by residual bootstrap and parametric bootstrap respectively. However, the nonparametric (residual) bootstrap leads to a more conservative conclusion (7-8 significant pairs), compared to the parametric one (11-13 significant pairs).

**Table 4.** Using residual (nonparametric) bootstrap methods: False coverage-statement rate (FCR) - Adjusted BH-Selected CIs for selected parameters indicated by the percentile bootstrap CI; All confidence intervals above show significance against *H*_0_ : *τ*_*i*_ = *τ*_*j*_.

| | $p$-value by percentile bootstrap CI | CI lower | CI upper | CI length | mean | sd | est |
| --- | --- | --- | --- | --- | --- | --- | --- |
| $\hat{\tau}_A^{bs} - \hat{\tau}_H^{bs}$ | $< 10^{-5}$ | -28.059 | -10.204 | 17.856 | -19.298 | 3.624 | -19.186 |
| $\hat{\tau}_B^{bs} - \hat{\tau}_H^{bs}$ | $< 10^{-5}$ | -25.265 | -8.512 | 16.753 | -17.707 | 3.213 | -18.115 |
| $\hat{\tau}_C^{bs} - \hat{\tau}_H^{bs}$ | $< 10^{-5}$ | -26.081 | -9.485 | 16.596 | -18.197 | 3.379 | -18.347 |
| $\hat{\tau}_D^{bs} - \hat{\tau}_F^{bs}$ | 0.008 | 0.659 | 17.297 | 16.638 | 8.125 | 3.348 | 7.544 |
| $\hat{\tau}_D^{bs} - \hat{\tau}_H^{bs}$ | $< 10^{-5}$ | -19.830 | -3.745 | 16.086 | -12.414 | 3.191 | -12.679 |
| $\hat{\tau}_E^{bs} - \hat{\tau}_H^{bs}$ | $< 10^{-5}$ | -24.668 | -7.003 | 17.665 | -15.496 | 3.489 | -15.233 |
| $\hat{\tau}_F^{bs} - \hat{\tau}_H^{bs}$ | $< 10^{-5}$ | -29.730 | -11.710 | 18.020 | -20.539 | 3.677 | -20.223 |
| $\hat{\tau}_G^{bs} - \hat{\tau}_H^{bs}$ | $< 10^{-5}$ | -24.095 | -7.635 | 16.460 | -16.482 | 3.187 | -16.687 |

**Table 5.** Using parametric bootstrap: False coverage-statement rate (FCR) - Adjusted BH-Selected CIs for selected parameters indicated by the percentile bootstrap CI; All confidence intervals above show significance against *H*_0_ : *τ*_*i*_ = *τ*_*j*_.

| | $p$-value by percentile bootstrap CI | CI lower | CI upper | CI length | mean | sd | est |
| --- | --- | --- | --- | --- | --- | --- | --- |
| $\hat{\tau}_A^{bs} - \hat{\tau}_D^{bs}$ | 0.002 | -10.778 | -3.105 | 7.674 | -6.578 | 1.545 | -6.507 |
| $\hat{\tau}_A^{bs} - \hat{\tau}_H^{bs}$ | $< 10^{-5}$ | -23.481 | -15.508 | 7.974 | -19.277 | 1.731 | -19.186 |
| $\hat{\tau}_B^{bs} - \hat{\tau}_D^{bs}$ | 0.006 | -9.552 | -1.525 | 8.026 | -5.301 | 1.525 | -5.436 |
| $\hat{\tau}_B^{bs} - \hat{\tau}_H^{bs}$ | $< 10^{-5}$ | -22.067 | -14.185 | 7.882 | -17.999 | 1.653 | -18.115 |
| $\hat{\tau}_C^{bs} - \hat{\tau}_D^{bs}$ | 0.006 | -10.075 | -1.356 | 8.719 | -5.662 | 1.644 | -5.668 |
| $\hat{\tau}_C^{bs} - \hat{\tau}_H^{bs}$ | $< 10^{-5}$ | -22.761 | -14.358 | 8.403 | -18.361 | 1.709 | -18.347 |
| $\hat{\tau}_D^{bs} - \hat{\tau}_F^{bs}$ | 0.002 | 2.768 | 12.825 | 10.057 | 7.680 | 1.828 | 7.544 |
| $\hat{\tau}_D^{bs} - \hat{\tau}_G^{bs}$ | 0.022 | 0.383 | 8.532 | 8.149 | 4.106 | 1.553 | 4.008 |
| $\hat{\tau}_D^{bs} - \hat{\tau}_H^{bs}$ | 0.002 | -16.687 | -8.478 | 8.209 | -12.698 | 1.765 | -12.679 |
| $\hat{\tau}_E^{bs} - \hat{\tau}_F^{bs}$ | 0.018 | 0.121 | 10.278 | 10.157 | 4.859 | 1.877 | 4.990 |
| $\hat{\tau}_E^{bs} - \hat{\tau}_H^{bs}$ | $< 10^{-5}$ | -20.403 | -11.297 | 9.106 | -15.519 | 1.925 | -15.233 |
| $\hat{\tau}_F^{bs} - \hat{\tau}_H^{bs}$ | $< 10^{-5}$ | -25.471 | -15.836 | 9.635 | -20.378 | 1.872 | -20.223 |
| $\hat{\tau}_G^{bs} - \hat{\tau}_H^{bs}$ | $< 10^{-5}$ | -20.893 | -12.667 | 8.225 | -16.804 | 1.648 | -16.687 |

#### Simultaneous Inference – maximum “unweighted” productivity *ψ*_*i*_

We now demonstrate that a wider variety of applications can be tackled by the proposed framework. For example, we can make inferences related to the maximum “unweighted” productivity. We consider using maximum “unweighted” productivity

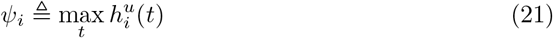

as the quantity of interest, where

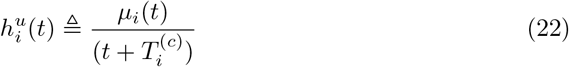

and 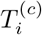 is the days of cultivation before the induction. The estimations of the “unweighted” productivity under different treatments are shown in Fig 6.

**Fig 6.**
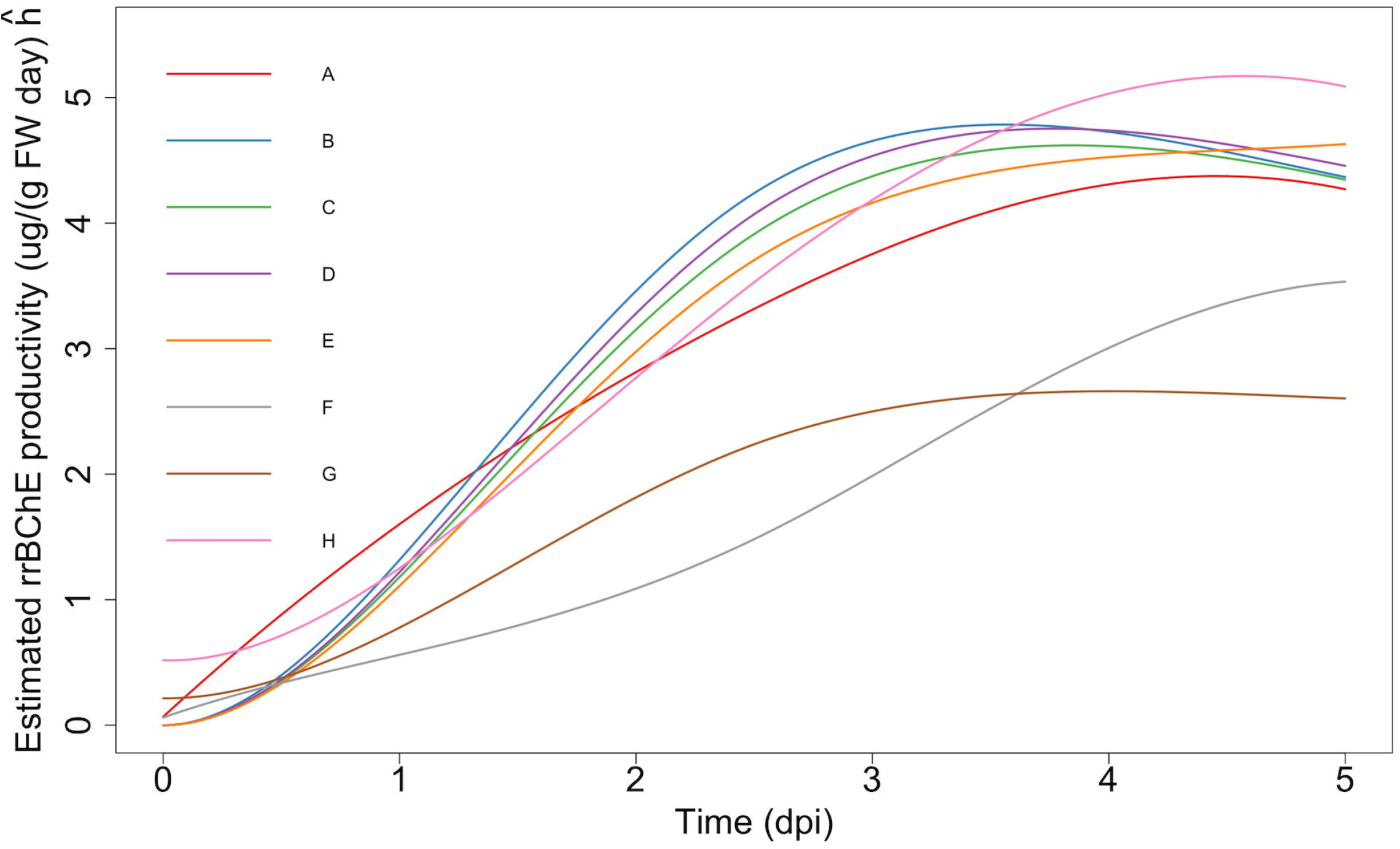
Estimated “unweighted” productivity curves based on observed data; Note that *T* ^(*c*)^ = (6, 6, 6, 7, 6, 8, 14, 8).

In this case, since the parameter is a nonlinear function of the trajectories, it is not possible to obtain the *p*-values using the null bootstrap distribution option. But we still do so by using the duality between hypothesis testing and finding confidence intervals.

We apply the same procedures we did for the maximum production level to the maximum “unweighted” productivity, using both parametric bootstrap and residual bootstrap methods. Table 6 and Table A3 in S1 Appendix show that residual bootstrap method leads to more conservative results (14 significant pairs), compared to the parametric one (21 significant pairs).

**Table 6.**
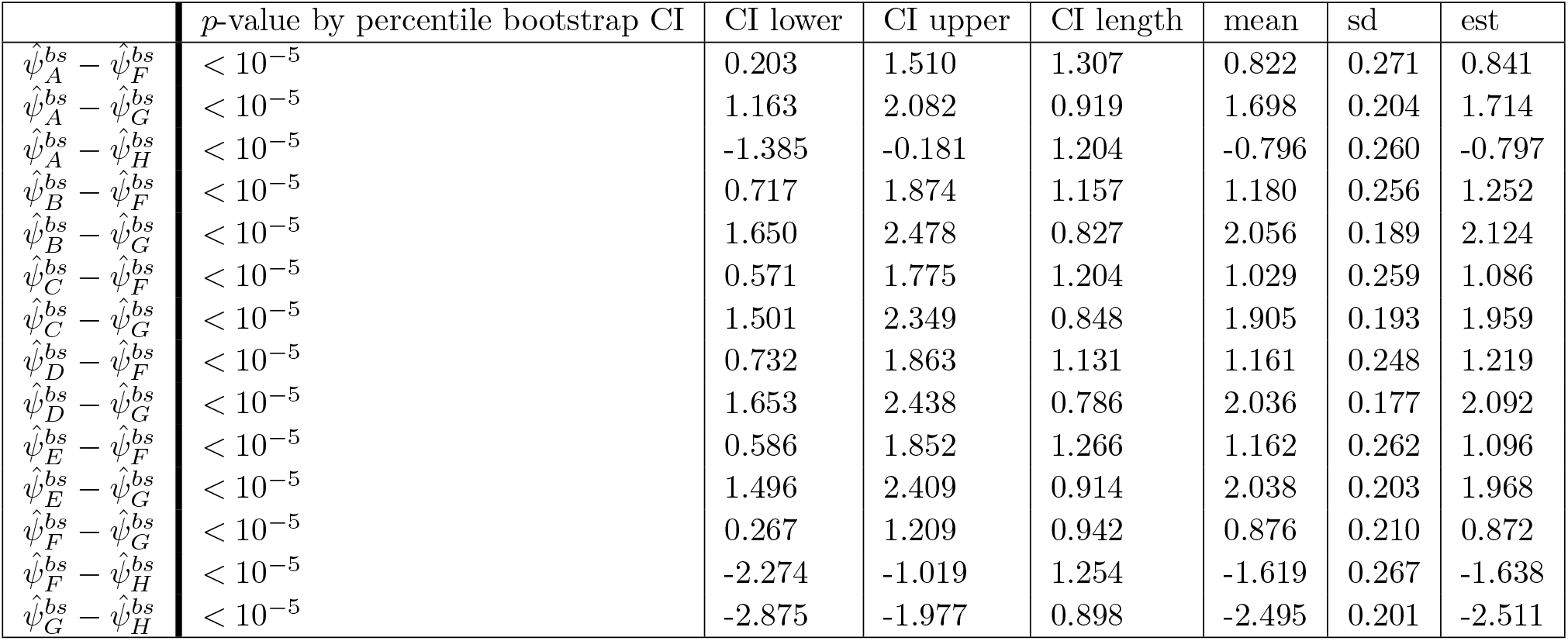
Using residual bootstrap method: False coverage-statement rate (FCR) - Adjusted BH-Selected CIs for selected parameters indicated by the percentile bootstrap CIs; All confidence intervals above show significance against *H*_0_ : *ψ*_*i*_ = *ψ*_*j*_.

### Summary of findings

#### Comparison between two versions of *p*-values

Given the results of two bootstrap methods, it turns out that the ranking of two versions of *p*-values are highly positively correlated (≥ 0.96), which means our method is robust and not sensitive to the way we compute *p*-values.

#### Comparison of residual bootstrap and parametric bootstrap

Table A1 and Table A2 in S1 Appendix may be compared for a comparative performance of different bootstrap strategies. First, within each table, we can see the *p*-values by using the null and duality of *p*-value and CI are consistent based on the rank and its correlation. Second, we can see that residual bootstrap is more conservative (Table A1 only has 7 significant differences) and *p*-values are larger. We can see the main difference depends on the variability of data (the way we generate bootstrap samples). Parametric bootstrap samples show higher variability (we make use of distribution assumption) while residual ones are with lower variability. This means the residual bootstrap is more conservative and the result is consistent with Fig 7.

**Fig 7.**
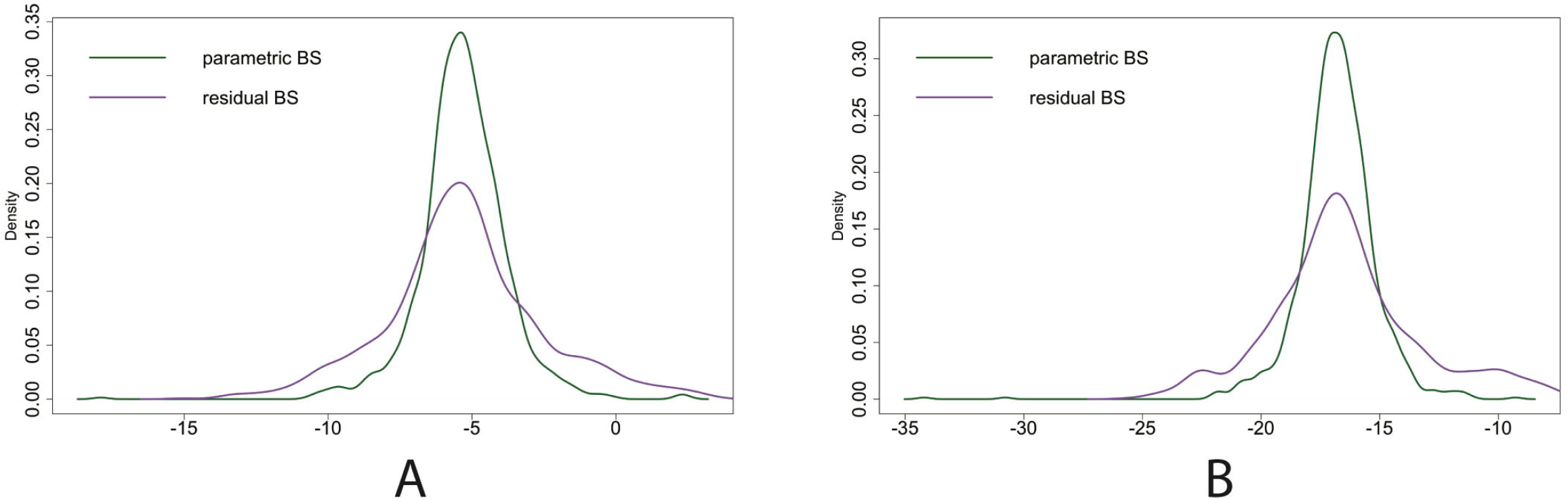
**Comparison: the empirical parametric bootstrap sampling distribution and the empirical residual bootstrap sampling distribution of 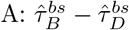 that is significant in parametric bootstrap case while not in residual bootstrap case; 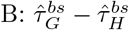 that is significant in both cases.**

Compared to the parametric bootstrap method, the residual bootstrap method is obviously more conservative for hypothesis testing. By comparing the parametric and residual bootstrap sampling distributions for parameters (see Fig 7), it is obvious that residual bootstrap method leads to more widely spread sampling distributions, which yields larger *p*-values and fewer significant testing results. Again, the residual bootstrap method is not particularly effective when the number of treatments is small. *I* = 8 might not be enough and a choice needs to be made to make inference.

## Discussion

### Implications for scientific investigation

The following conclusions can be made from our investigation. Our framework works well even when the data are limited so that standard asymptotic theory for inference is not applicable, which is very common in complicated biological experiments. In addition, our analysis is able to handle multiple questions of interest. Such an analysis can help suggest and validate the strategy of designing lengthy and expensive experiments, where the number of replicates is limited. We can see that the two variants of bootstrap methods work well even when the data are extremely limited. Nevertheless, it is suggested that one should always be cautious to make decisions on which type of bootstrap method to use. As far as we can see, if one is more confident in the model assumption then parametric bootstrap is the preferred option. Otherwise, the residual (nonparametric) bootstrap is a better choice as it is more conservative.

### Further extensions of the framework

The preceding analysis makes it clear that our framework is highly effective especially when multiple process metrics are of interest at the same time. In addition to the parameters of interest we mentioned, our framework allows us to extend the analysis to other parameters such as the average production and average “unweighted” productivity. The average production level corresponding to the *i*-th treatment can be described as 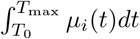 where *T*_0_ is the starting point. Corresponding null hypothesis is 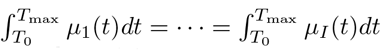. Similarly, the average “unweighted” productivity can be written as 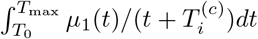 for the *i*-th treatment. Under the null we have the following equality 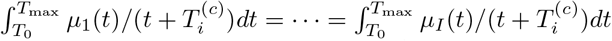. It should be noted that *p*-values cannot be calculated based on the null bootstrap distribution for average “unweighted” productivity because of nonlinearity of the trajectories. In this case, we adopt the percentile bootstrap option to obtain *p*-values for simultaneous inference.

## Supporting information

### S1 Appendix. Showing supplementary materials

Algorithm for double bootstrap procedure for computing 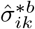, one figure and five tables are included.

