## Supplementary material for "Analysis of variability of functionals of recombinant protein production trajectories based on limited data": S1 Appendix

### Double bootstrap procedure for computing $\hat{\sigma}^{*b}_{ik}$

The procedure is as follows (assuming only one replicate):

1. Generate bootstrap samples $\{Y^{*b}_{ij}\}^{B}_{b=1}$, where $B$ is large (say $B = 1000$) and $Y^{*b}_{ij}$ is the $b$-th bootstrap measurement on $i$-th treatment at time $t_j$.

2. Let $\hat{\Theta}^{*b}_{i}$ be the corresponding bootstrap estimate of $\Theta_i$, and correspondingly, $\hat{\delta}^{*b}_{ik} = \hat{\Theta}^{*b}_{i} - \hat{\Theta}^{*b}_{k}$.

3. Estimate $\sigma^2_{ik}$ as follows:

$$\hat{\sigma}^2_{ik} = (B-1)^{-1} \sum_{b=1}^{B} (\hat{\delta}^{*b}_{ik} - \bar{\delta}^{*}_{ik})^2$$

   where $\bar{\delta}^{*}_{ik} = B^{-1} \sum_{b=1}^{B} \hat{\delta}^{*b}_{ik}$ is a bootstrap estimate of $E(\hat{\delta}_{ik})$.

4. In the percentile-$t$ bootstrap procedure, we compute a bootstrap estimate of $\hat{\sigma}_{ik}$, namely, $\hat{\sigma}^{*b}_{ik}$, corresponding to the $b$-th bootstrap sample used for inference. This will involve repeating the procedure as described above, but now treating the data for the $b$-th bootstrap sample ($Y^{*b}_{ij}$) as the raw data.

5. The procedure for computing $\hat{\sigma}^{*b}_{ik}$ is actually a so-called "double bootstrap" procedure. Again, these bootstrap samples for each $b$ will have to be generated independently. So, this is clearly a computationally intensive procedure.

### Figures

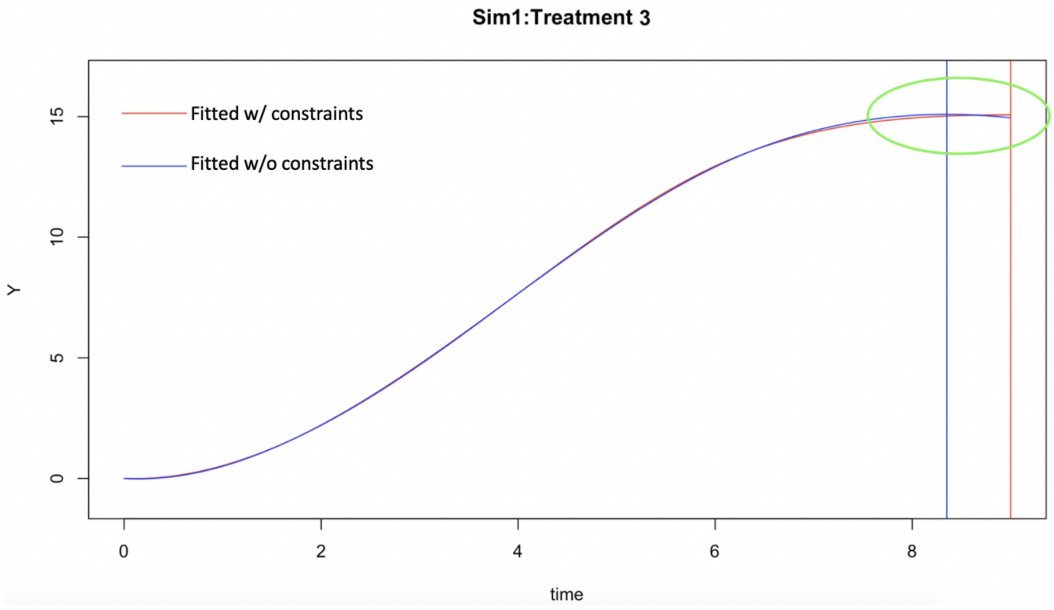


**Fig A1.** In the simulation study of Treatment 3: blue curve represents unconstrained estimation while the red curve is constrained estimation. Decreasing pattern shown at the end of time point in treatment 3; the peak value of unconstrained estimation is shown by blue vertical line while the red vertical line for the peak of constrained one.

### Tables

| | $p$-value based on the null bootstrap distribution | CI lower | CI upper | CI length | mean | sd | est |
|---|---|---|---|---|---|---|---|
| $\hat{\tau}_A^{bs} - \hat{\tau}_H^{bs}$ | $< 10^{-5}$ | -28.082 | -9.975 | 18.107 | -19.298 | 3.624 | -19.186 |
| $\hat{\tau}_B^{bs} - \hat{\tau}_H^{bs}$ | $< 10^{-5}$ | -25.337 | -8.495 | 16.842 | -17.707 | 3.213 | -18.115 |
| $\hat{\tau}_C^{bs} - \hat{\tau}_H^{bs}$ | $< 10^{-5}$ | -26.293 | -9.439 | 16.853 | -18.197 | 3.379 | -18.347 |
| $\hat{\tau}_D^{bs} - \hat{\tau}_H^{bs}$ | $< 10^{-5}$ | -20.150 | -3.602 | 16.547 | -12.414 | 3.191 | -12.679 |
| $\hat{\tau}_E^{bs} - \hat{\tau}_H^{bs}$ | $< 10^{-5}$ | -24.672 | -6.965 | 17.708 | -15.496 | 3.489 | -15.233 |
| $\hat{\tau}_F^{bs} - \hat{\tau}_H^{bs}$ | $< 10^{-5}$ | -29.974 | -11.644 | 18.330 | -20.539 | 3.677 | -20.223 |
| $\hat{\tau}_G^{bs} - \hat{\tau}_H^{bs}$ | $< 10^{-5}$ | -24.125 | -7.616 | 16.509 | -16.482 | 3.187 | -16.687 |

**Table A1.** Using **residual** (nonparametric) bootstrap methods: False coverage-statement rate (FCR) - Adjusted BH-Selected CIs for selected parameters indicated by **the null bootstrap distribution**; All confidence intervals above show significance against $H_0 : \tau_i = \tau_j$

| | $p$-value based on the null bootstrap distribution | CI lower | CI upper | CI length | mean | sd | est |
|---|---|---|---|---|---|---|---|
| $\hat{\tau}_A^{bs} - \hat{\tau}_D^{bs}$ | 0.003 | -10.996 | -3.011 | 7.985 | -6.578 | 1.545 | -6.507 |
| $\hat{\tau}_A^{bs} - \hat{\tau}_H^{bs}$ | $< 10^{-5}$ | -23.608 | -15.498 | 8.110 | -19.277 | 1.731 | -19.186 |
| $\hat{\tau}_B^{bs} - \hat{\tau}_D^{bs}$ | 0.004 | -9.687 | -1.330 | 8.358 | -5.301 | 1.525 | -5.436 |
| $\hat{\tau}_B^{bs} - \hat{\tau}_H^{bs}$ | $< 10^{-5}$ | -22.240 | -14.096 | 8.144 | -17.999 | 1.653 | -18.115 |
| $\hat{\tau}_C^{bs} - \hat{\tau}_D^{bs}$ | 0.011 | -10.215 | -1.116 | 9.099 | -5.662 | 1.644 | -5.668 |
| $\hat{\tau}_C^{bs} - \hat{\tau}_H^{bs}$ | $< 10^{-5}$ | -23.006 | -14.195 | 8.811 | -18.361 | 1.709 | -18.347 |
| $\hat{\tau}_D^{bs} - \hat{\tau}_F^{bs}$ | 0.005 | 2.673 | 12.901 | 10.228 | 7.680 | 1.828 | 7.544 |
| $\hat{\tau}_D^{bs} - \hat{\tau}_H^{bs}$ | $< 10^{-5}$ | -16.830 | -8.142 | 8.688 | -12.698 | 1.765 | -12.679 |
| $\hat{\tau}_E^{bs} - \hat{\tau}_H^{bs}$ | $< 10^{-5}$ | -20.601 | -11.252 | 9.349 | -15.519 | 1.925 | -15.233 |
| $\hat{\tau}_F^{bs} - \hat{\tau}_H^{bs}$ | $< 10^{-5}$ | -25.597 | -15.411 | 10.185 | -20.378 | 1.872 | -20.223 |
| $\hat{\tau}_G^{bs} - \hat{\tau}_H^{bs}$ | $< 10^{-5}$ | -21.068 | -12.309 | 8.759 | -16.804 | 1.648 | -16.687 |

**Table A2.** Using **parametric** bootstrap: False coverage-statement rate (FCR) - Adjusted BH-Selected CIs for selected parameters indicated by **the null bootstrap distribution**; All confidence intervals above show significance against $H_0 : \tau_i = \tau_j$

| | $p$-value based on the null bootstrap distribution | CI lower | CI upper | CI length | mean | sd | est |
|---|---|---|---|---|---|---|---|
| $\hat{\psi}_A^{bs} - \hat{\psi}_B^{bs}$ | 0.004 | -0.644 | -0.174 | 0.471 | -0.401 | 0.110 | -0.410 |
| $\hat{\psi}_A^{bs} - \hat{\psi}_D^{bs}$ | 0.008 | -0.594 | -0.158 | 0.436 | -0.369 | 0.106 | -0.378 |
| $\hat{\psi}_A^{bs} - \hat{\psi}_E^{bs}$ | 0.030 | -0.611 | -0.034 | 0.578 | -0.282 | 0.137 | -0.255 |
| $\hat{\psi}_A^{bs} - \hat{\psi}_F^{bs}$ | $< 10^{-5}$ | 0.507 | 1.127 | 0.620 | 0.823 | 0.135 | 0.841 |
| $\hat{\psi}_A^{bs} - \hat{\psi}_G^{bs}$ | $< 10^{-5}$ | 1.501 | 1.894 | 0.393 | 1.698 | 0.095 | 1.714 |
| $\hat{\psi}_A^{bs} - \hat{\psi}_H^{bs}$ | 0.002 | -1.055 | -0.568 | 0.486 | -0.809 | 0.123 | -0.797 |
| $\hat{\psi}_B^{bs} - \hat{\psi}_F^{bs}$ | $< 10^{-5}$ | 0.940 | 1.488 | 0.549 | 1.224 | 0.123 | 1.252 |
| $\hat{\psi}_B^{bs} - \hat{\psi}_G^{bs}$ | $< 10^{-5}$ | 1.914 | 2.261 | 0.347 | 2.099 | 0.081 | 2.124 |
| $\hat{\psi}_B^{bs} - \hat{\psi}_H^{bs}$ | 0.006 | -0.628 | -0.205 | 0.423 | -0.407 | 0.112 | -0.387 |
| $\hat{\psi}_C^{bs} - \hat{\psi}_F^{bs}$ | $< 10^{-5}$ | 0.763 | 1.312 | 0.549 | 1.055 | 0.123 | 1.086 |
| $\hat{\psi}_C^{bs} - \hat{\psi}_G^{bs}$ | $< 10^{-5}$ | 1.717 | 2.096 | 0.379 | 1.929 | 0.084 | 1.959 |
| $\hat{\psi}_C^{bs} - \hat{\psi}_H^{bs}$ | 0.002 | -0.813 | -0.374 | 0.439 | -0.577 | 0.112 | -0.552 |
| $\hat{\psi}_D^{bs} - \hat{\psi}_F^{bs}$ | $< 10^{-5}$ | 0.885 | 1.448 | 0.563 | 1.192 | 0.122 | 1.219 |
| $\hat{\psi}_D^{bs} - \hat{\psi}_G^{bs}$ | $< 10^{-5}$ | 1.885 | 2.212 | 0.327 | 2.067 | 0.078 | 2.092 |
| $\hat{\psi}_D^{bs} - \hat{\psi}_H^{bs}$ | 0.002 | -0.672 | -0.246 | 0.426 | -0.439 | 0.112 | -0.419 |
| $\hat{\psi}_E^{bs} - \hat{\psi}_F^{bs}$ | $< 10^{-5}$ | 0.800 | 1.426 | 0.626 | 1.105 | 0.143 | 1.096 |
| $\hat{\psi}_E^{bs} - \hat{\psi}_G^{bs}$ | $< 10^{-5}$ | 1.784 | 2.241 | 0.458 | 1.979 | 0.110 | 1.968 |
| $\hat{\psi}_E^{bs} - \hat{\psi}_H^{bs}$ | 0.008 | -0.778 | -0.236 | 0.542 | -0.527 | 0.138 | -0.542 |
| $\hat{\psi}_F^{bs} - \hat{\psi}_G^{bs}$ | $< 10^{-5}$ | 0.629 | 1.145 | 0.516 | 0.875 | 0.112 | 0.872 |
| $\hat{\psi}_F^{bs} - \hat{\psi}_H^{bs}$ | $< 10^{-5}$ | -1.906 | -1.342 | 0.564 | -1.631 | 0.135 | -1.638 |
| $\hat{\psi}_G^{bs} - \hat{\psi}_H^{bs}$ | $< 10^{-5}$ | -2.696 | -2.323 | 0.373 | -2.506 | 0.097 | -2.511 |

**Table A3.** Using **parametric** bootstrap method: False coverage-statement rate (FCR) - Adjusted BH-Selected CIs for selected parameters indicated by **the percentile bootstrap CIs**; All confidence intervals above show significance against $H_0 : \psi_i = \psi_j$

| | CI lower | CI upper | CI length | mean | sd | est |
|---|---|---|---|---|---|---|
| $\hat{\theta}_A^{bs}$ | 3.493 | 3.849 | 0.355 | 3.642 | 0.096 | 3.619 |
| $\hat{\theta}_B^{bs}$ | 2.758 | 2.978 | 0.220 | 2.841 | 0.064 | 2.813 |
| $\hat{\theta}_C^{bs}$ | 2.958 | 3.238 | 0.280 | 3.081 | 0.077 | 3.063 |
| $\hat{\theta}_D^{bs}$ | 2.513 | 2.678 | 0.165 | 2.587 | 0.046 | 2.588 |
| $\hat{\theta}_E^{bs}$ | 3.138 | 3.463 | 0.325 | 3.278 | 0.081 | 3.268 |
| $\hat{\theta}_F^{bs}$ | 4.174 | 4.484 | 0.310 | 4.318 | 0.077 | 4.314 |
| $\hat{\theta}_G^{bs}$ | 2.673 | 2.878 | 0.205 | 2.763 | 0.055 | 2.748 |
| $\hat{\theta}_H^{bs}$ | 2.578 | 2.743 | 0.165 | 2.659 | 0.042 | 2.658 |
| $\hat{\theta}_A^{bs} - \hat{\theta}_B^{bs}$ | 0.591 | 1.021 | 0.430 | 0.801 | 0.116 | 0.806 |
| $\hat{\theta}_A^{bs} - \hat{\theta}_C^{bs}$ | 0.315 | 0.801 | 0.485 | 0.561 | 0.122 | 0.556 |
| $\hat{\theta}_A^{bs} - \hat{\theta}_D^{bs}$ | 0.876 | 1.271 | 0.395 | 1.055 | 0.107 | 1.031 |
| $\hat{\theta}_A^{bs} - \hat{\theta}_E^{bs}$ | 0.130 | 0.631 | 0.501 | 0.364 | 0.125 | 0.350 |
| $\hat{\theta}_A^{bs} - \hat{\theta}_F^{bs}$ | -0.911 | -0.440 | 0.470 | -0.676 | 0.122 | -0.696 |
| $\hat{\theta}_A^{bs} - \hat{\theta}_G^{bs}$ | 0.686 | 1.086 | 0.400 | 0.879 | 0.110 | 0.871 |
| $\hat{\theta}_A^{bs} - \hat{\theta}_H^{bs}$ | 0.806 | 1.216 | 0.410 | 0.983 | 0.105 | 0.961 |
| $\hat{\theta}_B^{bs} - \hat{\theta}_C^{bs}$ | -0.430 | -0.060 | 0.370 | -0.240 | 0.102 | -0.250 |
| $\hat{\theta}_B^{bs} - \hat{\theta}_D^{bs}$ | 0.130 | 0.415 | 0.285 | 0.254 | 0.079 | 0.225 |
| $\hat{\theta}_B^{bs} - \hat{\theta}_E^{bs}$ | -0.631 | -0.245 | 0.385 | -0.437 | 0.103 | -0.455 |
| $\hat{\theta}_B^{bs} - \hat{\theta}_F^{bs}$ | -1.662 | -1.281 | 0.380 | -1.477 | 0.102 | -1.502 |
| $\hat{\theta}_B^{bs} - \hat{\theta}_G^{bs}$ | -0.075 | 0.240 | 0.315 | 0.078 | 0.087 | 0.065 |
| $\hat{\theta}_B^{bs} - \hat{\theta}_H^{bs}$ | 0.060 | 0.335 | 0.275 | 0.182 | 0.078 | 0.155 |
| $\hat{\theta}_C^{bs} - \hat{\theta}_D^{bs}$ | 0.335 | 0.691 | 0.355 | 0.494 | 0.089 | 0.475 |
| $\hat{\theta}_C^{bs} - \hat{\theta}_E^{bs}$ | -0.430 | 0.025 | 0.455 | -0.197 | 0.113 | -0.205 |
| $\hat{\theta}_C^{bs} - \hat{\theta}_F^{bs}$ | -1.441 | -1.006 | 0.435 | -1.237 | 0.109 | -1.251 |
| $\hat{\theta}_C^{bs} - \hat{\theta}_G^{bs}$ | 0.145 | 0.506 | 0.360 | 0.318 | 0.094 | 0.315 |
| $\hat{\theta}_C^{bs} - \hat{\theta}_H^{bs}$ | 0.260 | 0.621 | 0.360 | 0.422 | 0.089 | 0.405 |
| $\hat{\theta}_D^{bs} - \hat{\theta}_E^{bs}$ | -0.891 | -0.521 | 0.370 | -0.691 | 0.093 | -0.681 |
| $\hat{\theta}_D^{bs} - \hat{\theta}_F^{bs}$ | -1.907 | -1.552 | 0.355 | -1.731 | 0.089 | -1.727 |
| $\hat{\theta}_D^{bs} - \hat{\theta}_G^{bs}$ | -0.315 | -0.050 | 0.265 | -0.176 | 0.072 | -0.160 |
| $\hat{\theta}_D^{bs} - \hat{\theta}_H^{bs}$ | -0.190 | 0.045 | 0.235 | -0.072 | 0.063 | -0.070 |
| $\hat{\theta}_E^{bs} - \hat{\theta}_F^{bs}$ | -1.246 | -0.821 | 0.425 | -1.040 | 0.111 | -1.046 |
| $\hat{\theta}_E^{bs} - \hat{\theta}_G^{bs}$ | 0.330 | 0.721 | 0.390 | 0.515 | 0.101 | 0.521 |
| $\hat{\theta}_E^{bs} - \hat{\theta}_H^{bs}$ | 0.455 | 0.816 | 0.360 | 0.619 | 0.092 | 0.611 |
| $\hat{\theta}_F^{bs} - \hat{\theta}_G^{bs}$ | 1.376 | 1.732 | 0.355 | 1.555 | 0.094 | 1.567 |
| $\hat{\theta}_F^{bs} - \hat{\theta}_H^{bs}$ | 1.502 | 1.832 | 0.330 | 1.659 | 0.085 | 1.657 |
| $\hat{\theta}_G^{bs} - \hat{\theta}_H^{bs}$ | -0.020 | 0.245 | 0.265 | 0.104 | 0.068 | 0.090 |

**Table A4.** Parametric Bootstrap CIs for $c = 40$; Significance (at level $\alpha = 0.05$) against $H_0 : \theta_i = \theta_j$ is in red; differences involving treatment G or H (which we are most interested in) are highlighted in yellow; assuming $t$-distributed noise.

| | CI lower | CI upper | CI length | mean | sd | est |
|---|---|---|---|---|---|---|
| $\hat{\gamma}_A^{bs}$ | 4.194 | 5.000 | 0.806 | 4.491 | 0.183 | 4.424 |
| $\hat{\gamma}_B^{bs}$ | 3.569 | 3.834 | 0.265 | 3.634 | 0.110 | 3.594 |
| $\hat{\gamma}_C^{bs}$ | 3.634 | 4.274 | 0.641 | 3.882 | 0.185 | 3.859 |
| $\hat{\gamma}_D^{bs}$ | 3.589 | 3.939 | 0.350 | 3.722 | 0.117 | 3.719 |
| $\hat{\gamma}_E^{bs}$ | 4.089 | 5.000 | 0.911 | 4.761 | 0.325 | 5.000 |
| $\hat{\gamma}_F^{bs}$ | 4.750 | 5.000 | 0.250 | 4.954 | 0.074 | 5.000 |
| $\hat{\gamma}_G^{bs}$ | 3.539 | 3.899 | 0.360 | 3.647 | 0.162 | 3.599 |
| $\hat{\gamma}_H^{bs}$ | 4.374 | 4.965 | 0.591 | 4.514 | 0.123 | 4.459 |
| $\hat{\gamma}_A^{bs} - \hat{\gamma}_B^{bs}$ | 0.455 | 1.401 | 0.946 | 0.857 | 0.212 | 0.831 |
| $\hat{\gamma}_A^{bs} - \hat{\gamma}_C^{bs}$ | 0.090 | 1.196 | 1.106 | 0.609 | 0.255 | 0.566 |
| $\hat{\gamma}_A^{bs} - \hat{\gamma}_D^{bs}$ | 0.370 | 1.311 | 0.941 | 0.769 | 0.217 | 0.706 |
| $\hat{\gamma}_A^{bs} - \hat{\gamma}_E^{bs}$ | -0.771 | 0.551 | 1.321 | -0.270 | 0.373 | -0.576 |
| $\hat{\gamma}_A^{bs} - \hat{\gamma}_F^{bs}$ | -0.796 | 0.035 | 0.831 | -0.463 | 0.198 | -0.576 |
| $\hat{\gamma}_A^{bs} - \hat{\gamma}_G^{bs}$ | 0.480 | 1.401 | 0.921 | 0.844 | 0.241 | 0.826 |
| $\hat{\gamma}_A^{bs} - \hat{\gamma}_H^{bs}$ | -0.501 | 0.526 | 1.026 | -0.022 | 0.223 | -0.035 |
| $\hat{\gamma}_B^{bs} - \hat{\gamma}_C^{bs}$ | -0.621 | 0.065 | 0.686 | -0.248 | 0.217 | -0.265 |
| $\hat{\gamma}_B^{bs} - \hat{\gamma}_D^{bs}$ | -0.335 | 0.150 | 0.485 | -0.088 | 0.159 | -0.125 |
| $\hat{\gamma}_B^{bs} - \hat{\gamma}_E^{bs}$ | -1.426 | -0.395 | 1.031 | -1.127 | 0.340 | -1.406 |
| $\hat{\gamma}_B^{bs} - \hat{\gamma}_F^{bs}$ | -1.426 | -1.076 | 0.350 | -1.320 | 0.132 | -1.406 |
| $\hat{\gamma}_B^{bs} - \hat{\gamma}_G^{bs}$ | -0.265 | 0.220 | 0.485 | -0.013 | 0.196 | -0.005 |
| $\hat{\gamma}_B^{bs} - \hat{\gamma}_H^{bs}$ | -1.336 | -0.616 | 0.721 | -0.880 | 0.169 | -0.866 |
| $\hat{\gamma}_C^{bs} - \hat{\gamma}_D^{bs}$ | -0.165 | 0.541 | 0.706 | 0.160 | 0.217 | 0.140 |
| $\hat{\gamma}_C^{bs} - \hat{\gamma}_E^{bs}$ | -1.316 | 0.000 | 1.316 | -0.879 | 0.374 | -1.141 |
| $\hat{\gamma}_C^{bs} - \hat{\gamma}_F^{bs}$ | -1.346 | -0.666 | 0.681 | -1.072 | 0.198 | -1.141 |
| $\hat{\gamma}_C^{bs} - \hat{\gamma}_G^{bs}$ | -0.145 | 0.666 | 0.811 | 0.235 | 0.247 | 0.260 |
| $\hat{\gamma}_C^{bs} - \hat{\gamma}_H^{bs}$ | -1.076 | -0.225 | 0.851 | -0.632 | 0.224 | -0.601 |
| $\hat{\gamma}_D^{bs} - \hat{\gamma}_E^{bs}$ | -1.406 | -0.305 | 1.101 | -1.039 | 0.351 | -1.281 |
| $\hat{\gamma}_D^{bs} - \hat{\gamma}_F^{bs}$ | -1.406 | -0.951 | 0.455 | -1.232 | 0.139 | -1.281 |
| $\hat{\gamma}_D^{bs} - \hat{\gamma}_G^{bs}$ | -0.200 | 0.345 | 0.546 | 0.075 | 0.202 | 0.120 |
| $\hat{\gamma}_D^{bs} - \hat{\gamma}_H^{bs}$ | -1.206 | -0.506 | 0.701 | -0.792 | 0.170 | -0.741 |
| $\hat{\gamma}_E^{bs} - \hat{\gamma}_F^{bs}$ | -0.861 | 0.225 | 1.086 | -0.193 | 0.333 | 0.000 |
| $\hat{\gamma}_E^{bs} - \hat{\gamma}_G^{bs}$ | 0.385 | 1.456 | 1.071 | 1.114 | 0.362 | 1.401 |
| $\hat{\gamma}_E^{bs} - \hat{\gamma}_H^{bs}$ | -0.495 | 0.596 | 1.091 | 0.248 | 0.348 | 0.541 |
| $\hat{\gamma}_F^{bs} - \hat{\gamma}_G^{bs}$ | 1.006 | 1.456 | 0.450 | 1.307 | 0.176 | 1.401 |
| $\hat{\gamma}_F^{bs} - \hat{\gamma}_H^{bs}$ | 0.000 | 0.606 | 0.606 | 0.440 | 0.141 | 0.541 |
| $\hat{\gamma}_G^{bs} - \hat{\gamma}_H^{bs}$ | -1.326 | -0.551 | 0.776 | -0.866 | 0.207 | -0.861 |

**Table A5.** Parametric Bootstrap CIs for $c = 0.1$; Significance (at level $\alpha = 0.05$) against $H_0 : \gamma_i = \gamma_j$ is in red; differences involving treatment G or H (which we are most interested in) are highlighted in yellow; assuming $t$-distributed noise.
